# Suppression of store-operated calcium entry causes dilated cardiomyopathy of the *Drosophila* heart

**DOI:** 10.1101/659136

**Authors:** Courtney E. Petersen, Matthew J. Wolf, Jeremy T. Smyth

## Abstract

Store-operated Ca^2+^ entry (SOCE) is an essential Ca^2+^ signaling and homeostatic mechanism present in nearly all animal cells. SOCE refers to influx of Ca^2+^ into cells that is activated by depletion of endoplasmic or sarcoplasmic reticulum stores (ER/SR) Ca^2+^ stores. In the SOCE pathway, STIM proteins function as Ca^2+^ sensors in the ER, and upon ER Ca^2+^ store depletion STIM rearranges to ER-plasma membrane junctions where it activates Orai Ca^2+^ influx channels. Multiple studies have implicated STIM and Orai mediated SOCE in the pathogenesis of cardiac hypertrophy. Importantly however, the functional roles of SOCE in normal heart physiology have not been well defined. We have addressed this in *Drosophila melanogaster*, a powerful animal model of cardiac development and physiology. We show that heart specific suppression of *Drosophila Stim* and *Orai* resulted in reduced contractility consistent with dilated cardiomyopathy, characterized by increased end diastolic and end systolic dimensions and decreased fractional shortening. Reduced contractility was apparent in larval hearts and became more pronounced in adults. Myofibers were disorganized and more widely spaced in larval and adult hearts with *Stim* and *Orai* RNAi as compared to controls, possibly reflecting decompensation or upregulated stress response signaling due to altered Ca^2+^ homeostasis. Lastly, we show that reduced heart function significantly affected animal health and viability, as animals with heart specific *Stim* and *Orai* suppression exhibited significant delays in post-embryonic development and adults died significantly earlier than controls. Collectively, our results demonstrate that SOCE is essential for normal heart physiology and establish *Drosophila* as an important model for delineation of functional SOCE roles in cardiomyocytes.

## INTRODUCTION

Cardiac disease continues to be a leading cause of morbidity and mortality throughout the western world, with current treatment options limited to palliative pharmacological or invasive therapy (McKenna et al., 2017). The discovery of curative treatments depends on thorough understanding of the molecular mechanisms that govern the onset and progression of cardiac pathophysiology. Significantly, irregularities in cardiomyocyte calcium (Ca^2+^) homeostasis are a major contributing factor to cardiac disease pathogenesis, and targeting Ca^2+^ signaling mechanisms may therefore be an important approach to novel therapeutic development (Abraham and Wolf, 2013; Kranias and Bers, 2007; Limas et al., 1987; MacLeod, 2016).

The role of Ca^2+^ in the process of excitation-contraction (E-C) coupling, which drives cardiomyocyte contractility, is well established. In E-C coupling, membrane depolarization opens L-type voltage gated Ca^2+^ channels and entering Ca^2+^ then activates ryanodine receptors (RyRs) in the sarcoplasmic reticulum (SR). Release of SR Ca^2+^ via RyRs results in a large cytoplasmic Ca^2+^ pulse that drives acto-myosin contractility (Bers, 2002). Ca^2+^ is also an important regulator of cardiomyocyte signaling pathways, such as those that control differentiation, cell growth, and pathological remodeling (Vega et al., 2003). Maintenance of Ca^2+^ homeostasis, including SR Ca^2+^ stores, is therefore essential to multiple aspects of cardiomyocyte physiology. Store-operated Ca^2+^ entry (SOCE) is a process that plays a major role in maintaining cellular Ca^2+^ homeostasis, as it couples the influx of extracellular Ca^2+^ to the depletion of sarco/endoplasmic (S/ER) Ca^2+^ stores. Ca^2+^ that enters the cell via SOCE can be pumped back into the S/ER to replenish depleted stores and restore S/ER Ca^2+^ homeostasis. Importantly, despite the prominent role for SR Ca^2+^ stores in cardiomyocyte physiology, the functions of SOCE in cardiomyocytes and overall cardiac physiology are poorly understood.

The main components of the SOCE pathway are STIM (stromal interacting molecule) and Orai. STIM is a single-pass transmembrane protein that serves as an E/SR Ca^2+^ sensor via its N-terminal. E/SR lumenal Ca^2+^ binding EF-hand domain, and Orai is a SOCE pore forming channel subunit in the plasma membrane (Putney, 2011, 2018). In response to E/SR Ca^2+^ store depletion, STIM undergoes a large conformational change that results in oligomerization and exposure of a cytoplasmic Orai activating domain. Oligomerized STIM then translocates to E/SR-plasma membrane junctions where it interacts with and activates Orai to induce Ca^2+^ influx (Putney, 2018; Smyth et al., 2010). In mammals, there are two STIM isoforms, STIM1 and STIM2, and three Orai isoforms (Orai1-3), with STIM1 and Orai1 exhibiting the widest functional distribution across mammalian cell and tissue types.

Numerous studies strongly suggest that SOCE contributes to the pathogenesis of pathological cardiac hypertrophy, whereby heart muscle mass increases in response to stressors such as hypertension or valve dysfunctions. For example, induction of cardiac hypertrophy by pressure overload results in upregulation of STIM1 and Orai1 expression in mouse cardiac tissue, and cardiomyocyte-specific STIM1 and Orai1 suppression attenuates the hypertrophic response (Benard et al., 2016; Hulot et al., 2011; Luo et al., 2012; Parks et al., 2016). Similarly, suppression of STIM1 or Orai1 in rodent cardiomyocytes attenuates phenylephrine and endothelin-1 induced hypertrophy (Hulot et al., 2011; Luo et al., 2012; Voelkers et al., 2010). Enhanced SOCE in response to pathological stimuli likely activates the calcineurin-nuclear factor of activated T-cells (NFAT) signaling axis, which is essential for reactivation of developmental gene expression and promotion of cardiomyocyte growth (Hulot et al., 2011; Molkentin et al., 1998; Schulz and Yutzey, 2004; Wilkins et al., 2004).

In light of this strong evidence that enhanced SOCE can drive pathological responses in cardiomyocytes, an important question remains: what is the role of SOCE in healthy cardiomyocytes and normal heart physiology? In support of the concept that SOCE is essential for healthy cardiac function, two independent studies have shown that cardiomyocyte restricted STIM1 deletion in mice results in marked left ventricular dilation and reduced ejection fraction in adult hearts (Collins et al., 2014; Horton et al., 2014; Parks et al., 2016). Decreased cardiac function was concomitant with indications of ER stress and changes in cardiomyocyte mitochondrial morphology (Collins et al., 2014), as well as altered contractile Ca^2+^ transients and myofibril organization (Parks et al., 2016). In addition, Orai1 suppression in zebrafish embryos resulted in reduced fractional shortening and severe heart failure (Volkers et al., 2012). These results support the conclusion that SOCE is essential for normal cardiac physiology, but importantly, the specific cellular processes that are regulated by SOCE in cardiomyocytes remain unknown. It is also unclear whether these results reflect full suppression of SOCE activity, because functional contributions by other STIM and Orai isoforms cannot be ruled out in these vertebrate models. The goal of our current study was to begin addressing these important questions by testing the role of SOCE in *Drosophila melanogaster* heart function, a valuable animal model in which powerful genetic tools can be integrated with *in vivo* analyses of cardiomyocyte physiology and heart function.

The *Drosophila* heart is a muscular tube of cardiomyocytes that runs along the dorsal midline of the animal. It is responsible for pumping hemolymph, a plasma-like fluid, throughout the body in an open circulatory system (Rotstein and Paululat, 2016). Importantly, the contractile physiology of the *Drosophila* heart, including cardiomyocyte Ca^2+^ transport mechanisms and sarcomere composition, is highly conserved with mammals (Lin et al., 2011; Ocorr et al., 2007), and the genetic and functional bases of many cardiomyopathies can be readily modeled and analyzed in flies (Piazza and Wessells, 2011). Simplified genetics is another important advantage of *Drosophila* over other animal models. In particular, *Drosophila* express single isoforms of *Stim* and *Orai*, thus precluding complications of functional overlap encountered with vertebrate models. Taking advantage of the strengths of the *Drosophila* model, we demonstrate that animals with heart-specific suppression of the key SOCE pathway components, *Stim* and *Orai*, exhibit dilated cardiomyopathy characterized by enlarged end-diastolic and end-systolic dimensions and decreased fractional shortening. Myofibrils were also disorganized and loosely spaced in *Stim* and *Orai* suppressed hearts, further consistent with disrupted contractile physiology. Lastly, these animals exhibited significantly delayed development and they died earlier than controls, suggesting pathological impairment of cardiac function. Our results, as well as those from other animal models, demonstrate that SOCE has essential and highly conserved roles in supporting normal heart physiology, and lay the groundwork for future studies using *Drosophila* to mechanistically define SOCE functions in the heart.

## RESULTS

### *Stim* and *Orai* suppression results in dilated cardiomyopathyx

*Drosophila Stim* and *Orai* loss-of-function mutants fail to grow properly and die as second or third instar larvae (Pathak et al., 2017), limiting their use in analysis of heart function. We therefore used inducible RNAi to suppress *Stim* and *Orai* expression specifically in the heart and avoid the systemic effects of global SOCE suppression. We first tested the effectiveness of our RNAi constructs by expressing them ubiquitously in the whole animal and analyzing transcript levels and phenotypes. *Stim* and *Orai* RNAi driven by the ubiquitous *act-GAL4* driver suppressed *Stim* and *Orai* mRNA expression in whole first instar larvae by 72.67 ± 3.18% and 80 ± 2.65% (mean ± SEM), respectively, compared to non-targeting RNAi controls, demonstrating potent knockdown of the targeted transcripts (Supplemental Figure S1A). Ubiquitous expression of *Stim* and *Orai* RNAi also resulted in reduced growth and larval lethality (Supplemental Figure S1B-F) similar to loss-of-function mutants, suggesting specific knockdown of the targeted gene products with few to no off-target effects.

We next used optical coherence tomography (OCT) to evaluate heart contractility in fully intact, non-anesthetized animals with heart specific *Stim* and *Orai* suppression. Adult males with heart-specific *tinC-GAL4* driven *Stim* and *Orai* RNAi exhibited significantly increased end-diastolic dimensions (EDD), and a more pronounced increase in end-systolic dimensions (ESD) as compared to non-targeting RNAi controls (Figure 1A-H).This resulted in significantly reduced fractional shortening (FS), a direct measure of the contractile strength of the heart, in *Stim* and *Orai* RNAi animals (88.73 ± 2.04% in control, 42.28 ± 1.4% in *Stim* RNAi, 49.21 ± 3.11% in *Orai RNAi*; mean ± SEM, p < 0.001) (Figure 1I). Enlarged ESD and EDD and reduced FS in *Stim* and *Orai* RNAi animals are consistent with dilated cardiomyopathy. Similar results were also observed using a second heart specific driver, *4xhand-GAL4* (Figure 1J-R). Animals with *4xhand-GAL4* driven *Stim* and *Orai* RNAi again exhibited significantly larger EDD and ESD compared to nontargeting RNAi controls (Figure 1P,Q), and this resulted in significantly reduced FS in *Orai* RNAi animals (Figure 1R). The decrease in FS was not significantly different in animals with *4xhand-GAL4* driven *Stim* RNAi, likely due to less efficient knockdown of *Stim* compared to *Orai* expression (see Supplemental Figure S1A). Neither *tinC-GAL4* nor *4xhand-GAL4* driven *Stim* and *Orai* RNAi significantly altered heart rates or caused notable arrhythmias compared to controls (Supplemental Figure S2A and B). Overall, heart specific suppression of *Stim* and *Orai* driven by two independent heart specific drivers resulted in dilated cardiomyopathy, suggesting an essential function for SOCE in adult *Drosophila* cardiac function.

**Figure 1:**
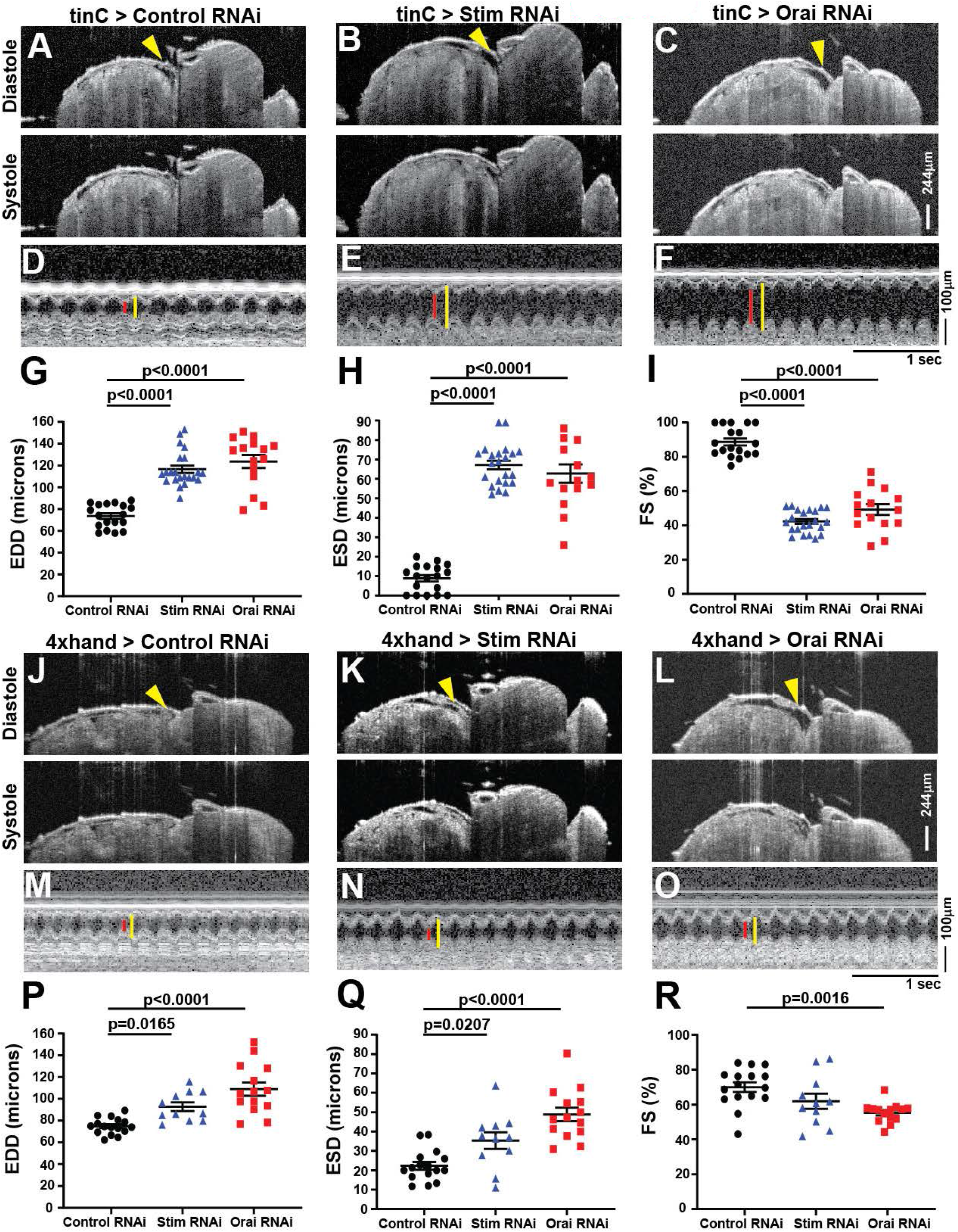
Heart specific *Stim* and *Orai* suppression results in dilated cardiomyopathy in adults. **A-C.** Representative OCT longitudinal B-mode images of five-day old adult male flies with *tinC-GAL4* driven non-targeting control, *Stim*, and *Orai* RNAi during diastole (upper panels) and systole (lower panels). Arrowheads indicate the heart in diastole images. **D-F.** Representative M-mode recordings from *tinC-GAL4* driven non-targeting control, *Stim*, and *Orai* RNAi with red lines depicting systole and yellow lines diastole. EDD (**G**), ESD (**H**), and FS (**I**) were calculated from M-mode recordings of five-day old males, and each symbol represents a measurement from a single animal. Bars indicate mean ± SEM, and p-values were calculated from One-way ANOVA with Tukey’s Multiple Comparison. **J-L.** Representative OCT longitudinal B-mode images of five-day old adult male flies with *4xhand-GAL4* driven non-targeting control, *Stim*, and *Orai* RNAi during diastole (upper panels) and systole (lower panels). Arrowheads indicate the heart in diastole images. **M-O.** Representative M-mode recordings from *4xhand-GAL4* driven non-targeting control, *Stim*, and *Orai* RNAi with red lines depicting systole and yellow lines diastole. EDD (**P**), ESD (**Q**), and FS (**R**) were calculated from M-mode recordings of five-day old males, and each symbol represents a measurement from a single animal. Bars indicate mean ± SEM, and p-values were calculated from One-way ANOVA with Tukey’s Multiple Comparison.

Results from mouse models demonstrate that STIM1 suppression results in age-dependent cardiomyopathy and heart failure in adults (Collins et al., 2014; Parks et al., 2016), whereas severe heart failure was evident during embryogenesis in zebrafish with Orai1 suppression (Volkers et al., 2012). We therefore sought to determine whether cardiomyopathy caused by SOCE suppression in *Drosophila* occurs in larvae, prior to the adult life cycle stage. Larval heart contractility was analyzed using intravital fluorescence imaging of animals with cardiomyocyte-specific tdTomato (CM-tdTom) expression (Klassen et al., 2017) as opposed to OCT, because OCT imaging in larvae is complicated by light scattering caused by high fat content. We first validated our intravital CM-tdTom imaging approach by showing that analysis of adult heart contractility yielded similar results to OCT, with significantly larger ESD and EDD and reduced FS in *tinC-GAL4* driven *Stim* RNAi hearts compared to controls (Figure S3, Supplemental Videos 1 and 2). Similar to adults, larvae with *tinC-GAL4* driven *Stim* and *Orai* RNAi exhibited significantly increased EDD and ESD compared to nontargeting RNAi controls (Figure 2A-H, Supplemental Videos 3 and 4), and this resulted in significantly decreased FS for *Orai*, though not *Stim* RNAi (Figure 2I).We also observed a small though significant decrease in heart rate in *Orai* but not *Stim* RNAi larvae (Supplemental Figure S2C). These results suggest that reduced cardiac output due to SOCE suppression is already evident in larval stages of *Drosophila* development, and thus is not exclusively an age-dependent effect in the adult heart. We also tested larval contractility with *4xhand-GAL4* driven *Stim* and *Orai* RNAi and although this resulted in modest increases in ESD and EDD, FS was unchanged compared to controls (Figure 2J-R). It is possible that *4xhand-GAL4* does not drive RNAi expression as strongly in the larval heart as does *tinC-GAL4*, thus explaining the weaker effects of *4xhand-GAL4*-driven *Stim* and *Orai* RNAi on larval heart contractility. Because of this, the *4xhand-GAL4* driver was not used further for larval heart analysis.

**Figure 2:**
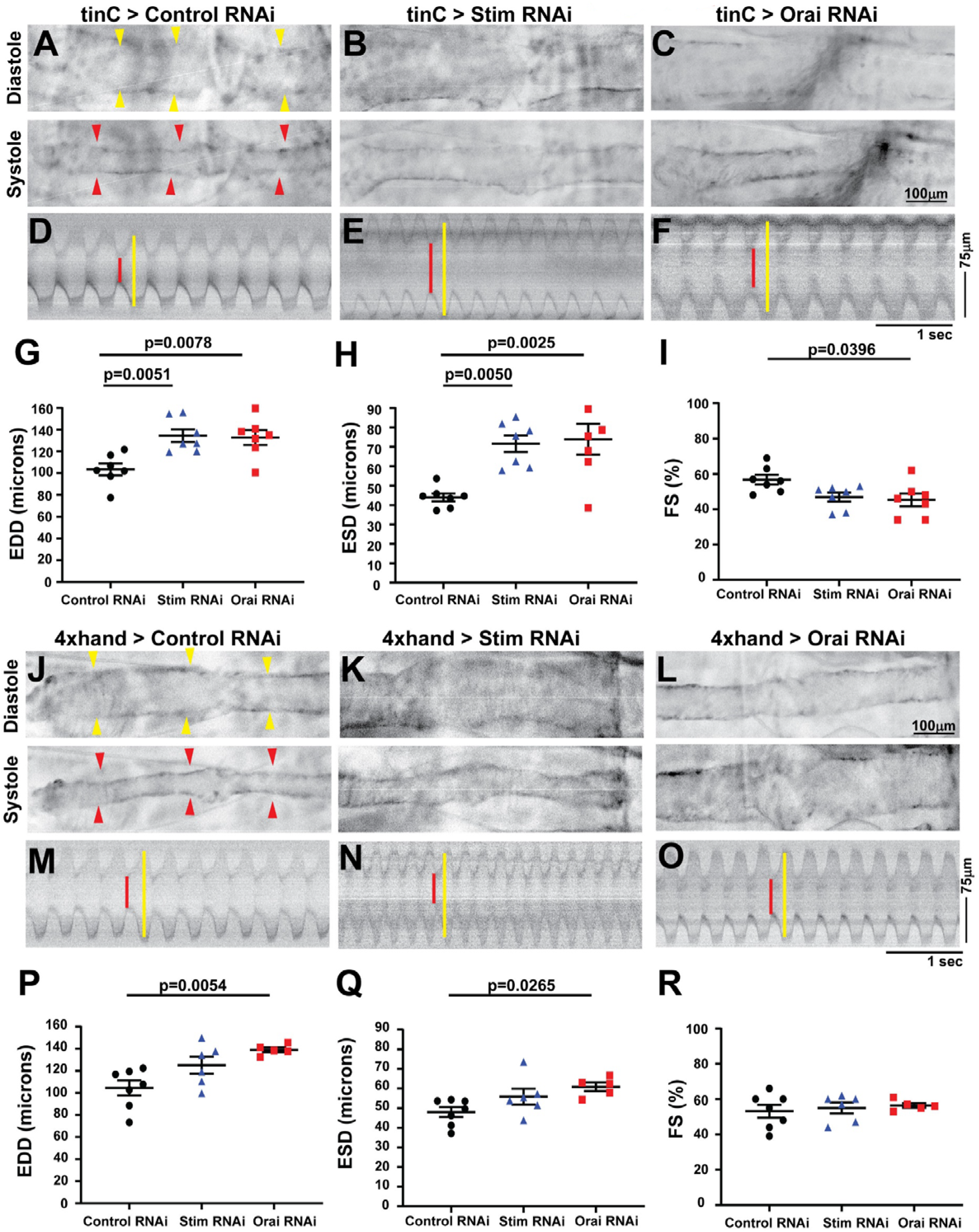
Larval *Stim* and *Orai* suppressed hearts are significantly dilated. **A-C** Representative longitudinal images from intravital fluorescence imaging of R94C02-tdTom expressing third instar larval hearts with *tinC-GAL4* driven non-targeting control, *Stim*, and *Orai* RNAi during diastole (upper panels) and systole (lower panels). Images are presented in inverted grayscale for added clarity, due to low fluorescence intensity. Yellow arrowheads indicate heart in diastolic images and red arrowheads in systolic images. **D-F.** Representative M-mode images from *tinC-GAL4* driven non-targeting control, *Stim*, and *Orai* RNAi hearts. Red lines depict systole and yellow lines diastole. EDD (**G**), ESD (**H**), FS (**I**) were calculated from M-mode recordings of third instar larva, and each symbol represents a measurement from a single animal. Bars indicate mean ± SEM, and p-values were calculated from One-way ANOVA with Tukey’s Multiple Comparison. **J-L.** Representative longitudinal images from intravital fluorescence imaging of R94C02-tdTom expressing third instar larval hearts with *4xhand-GAL4* driven non-targeting control, *Stim*, and *Orai* RNAi during diastole (upper panels) and systole (lower panels). Yellow arrowheads indicate heart in diastolic images and red arrowheads in systolic images. **M-O.** Representative M-mode images from *4xhand-GAL4* driven non-targeting control, *Stim*, and *Orai* RNAi hearts. Red lines depict systole and yellow lines diastole. EDD (**P**), ESD (**Q**), FS (**R**) were calculated from M-mode recordings of third instar larvae, and each symbol represents a measurement from a single animal. Bars indicate mean ± SEM, and p-values were calculated from One-way ANOVA with Tukey’s Multiple Comparison.

### Stim and Orai suppression disrupts myofibril organization

Disrupted myofibril organization is a common feature of dilated cardiomyopathy in mammalian (Garfinkel et al., 2018; Sussman et al., 1998) and *Drosophila* (Wolf et al., 2006) hearts. We therefore determined whether SOCE suppressed hearts similarly exhibit disorganized myofibrils. As shown in Figure 3A and B, hearts from *tinC-GAL4* driven control RNAi third-instar larvae exhibited evenly spaced myofibrils that were oriented circularly around the heart tube as revealed by actin staining and confocal microscopy. In contrast, myofibrils in *tinC-GAL4* driven *Stim* and *Orai* RNAi hearts exhibited highly varied orientations, with some myofibrils running nearly parallel to the long axis of the heart tube (Figure 3C-D). In addition, myofibrils in *Stim* and *Orai* RNAi hearts were more widely spaced compared to controls, as indicated by a significant reduction of nearly 45% in myofibril density in *Stim* and *Orai* RNAi versus control hearts (Figure 3E). In adult control animals, myofibrils again were densely packed and uniformly oriented around the heart tube (Figure 4A-B and E). Conversely, myofibrils in adult *tinC-GAL4* driven *Stim* and *Orai* RNAi hearts exhibited non-uniform orientation including longitudinal fibers, as well as significantly reduced myofibril density compared to controls (Figure 4C-D and H). Similar results were seen in adult hearts with *4xhand-GAL4* driven RNAi (Figure 4F-G), including a significant decrease in myofibril density (Figure I). These results are consistent with the dilated cardiomyopathy phenotype in SOCE suppressed animals, and suggest that myofibril remodeling as a result of suppressed SOCE may contribute to the aberrant contractility.

**Figure 3:**
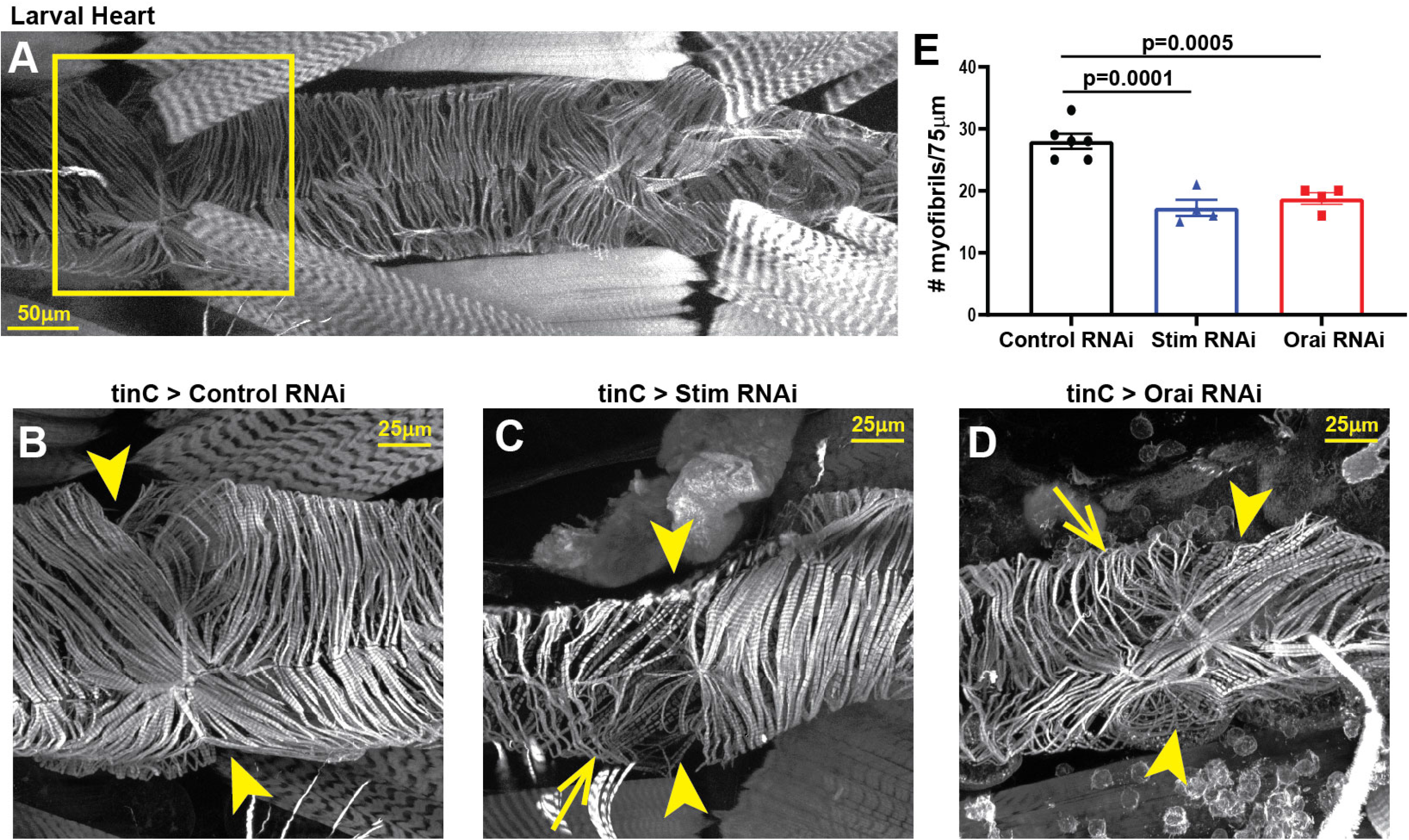
*Stim* and *Orai* suppression results in myofibril disorganization in larval hearts. **A.** Representative image of a control third instar larval heart stained with Phalloidin to visualize actin. The box denotes the region shown in the high resolution images. **B-D.** Representative high resolution images of the region around the second ostium of third instar larval hearts with *tinC-GAL4* driven non-targeting control, *Stim,* and *Orai* RNAi. Arrowheads point to ostia, and arrows point to regions with disorganized myofibrils. **E.** Plot of the total number of myofibrils per 75μm in larval hearts with *tinC-GAL4* driven non-targeting control, *Stim* and *Orai* RNAi. Each symbol represents a measurement from a single heart, and data are mean ± SEM (p-values were calculated from One-way ANOVA with Tukey’s Multiple Comparisons Test).

**Figure 4:**
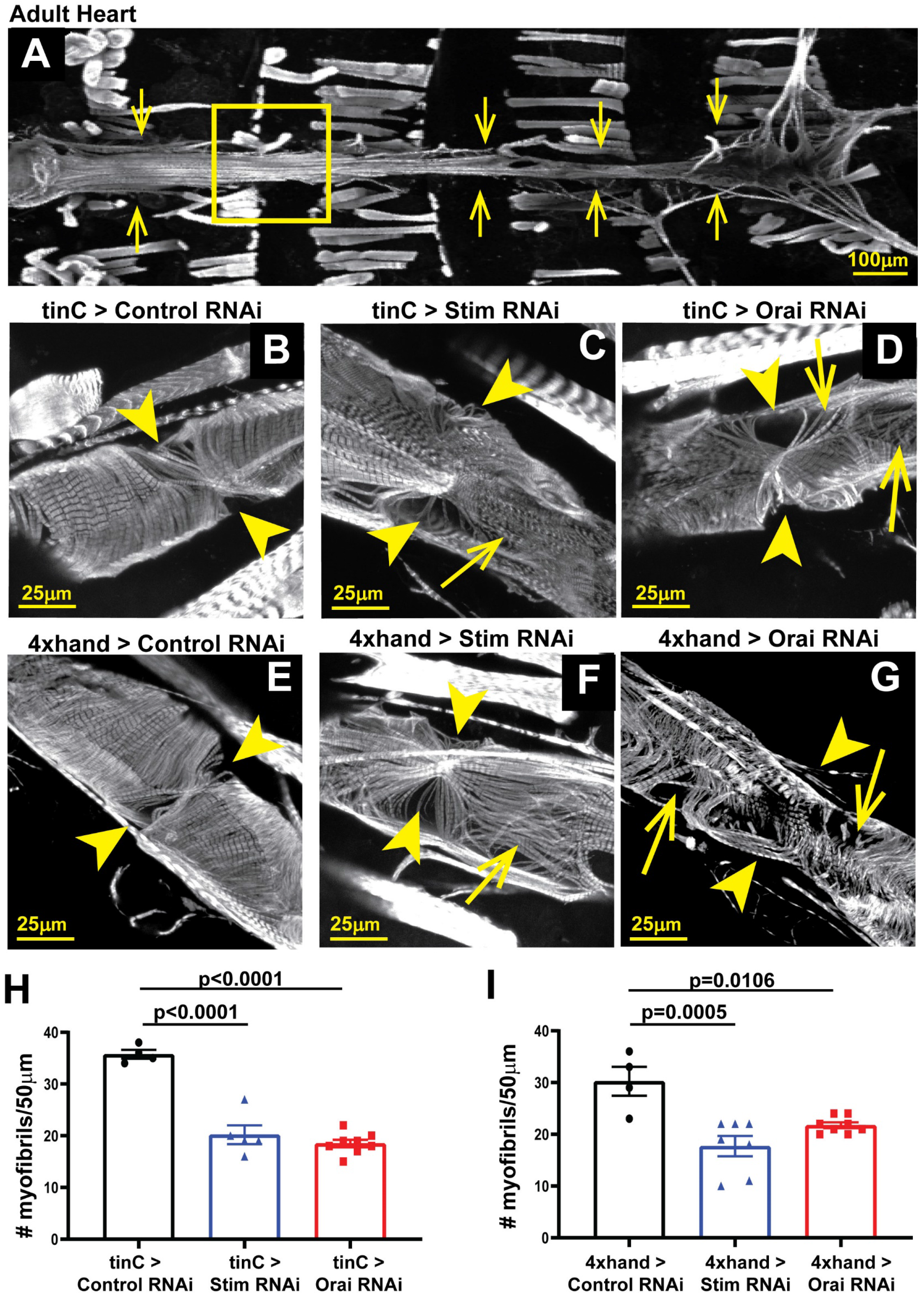
*Stim* and *Orai* suppression results in myofibril disorganization in adult hearts. **A.** Representative image of an adult control heart (indicated by arrows) stained with Phalloidin to visualize actin. The box denotes the region shown in the high-resolution images. **B-D.** Representative high resolution images of the region around the third ostium of five-day old adult male hearts with *tinC-GAL4* driven non-targeting control, *Stim*, and *Orai* RNAi. Arrowheads point to ostia, and arrows point to regions with disorganized myofibrils. **E-G.** Representative high-resolution images of the region around the third ostium of five-day old adult male hearts with *4xhand-GAL4* driven non-targeting control, *Stim*, and *Orai* RNAi. Arrowheads point to ostia, and arrows point to regions with disorganized myofibrils. **H.** Plot of the total number of myofibrils per 50μm in adult hearts with *tinC-GAL4* driven non-targeting control, *Stim* and *Orai* RNAi. Each symbol represents a measurement from a single heart, and data are mean ± SEM (p-values were calculated from One-way ANOVA with Tukey’s Multiple Comparisons Test). **I.** Plot of the total number of myofibrils per 50μm in adult hearts with *4xhand-GAL4* driven non-targeting control, *Stim* and *Orai* RNAi. Each symbol represents a measurement from a single heart, and data are mean ± SEM (p-values were calculated from One-way ANOVA with Tukey’s Multiple Comparisons Test).

### Heart specific *Stim* and *Orai* suppression reduces animal viability

We next determined whether the functional defects in *Stim* and *Orai* suppressed hearts affect animal health and viability. In support of this, larval development was significantly prolonged in animals with *tinC-GAL4* driven *Stim* and *Orai* RNAi, with nearly 86% fewer *Stim* and *Orai* RNAi animals having undergone pupariation on day five post-embryogenesis compared to controls (Figure 5A, B). Numbers of pupariated *Stim* and *Orai* RNAi animals were indistinguishable from controls by day six post-embryogenesis, indicating that the overall delay in pupariation was approximately 24 hours. Eclosion, when adult animals emerge from their pupal cases, was also delayed by about 24 hours in *tinC-GAL4* driven *Stim* and *Orai* animals compared to controls, suggesting that development was not further prolonged during pupal metamorphosis (Figure 2C-D). These results suggest that the contractile deficits caused by *Stim* and *Orai* suppression in the larval heart alter developmental physiology of the animal, possibly by limiting nutrient availability to developing tissues or reducing the circulation of the hormone ecdysone that regulates larval developmental timing.

**Figure 5:**
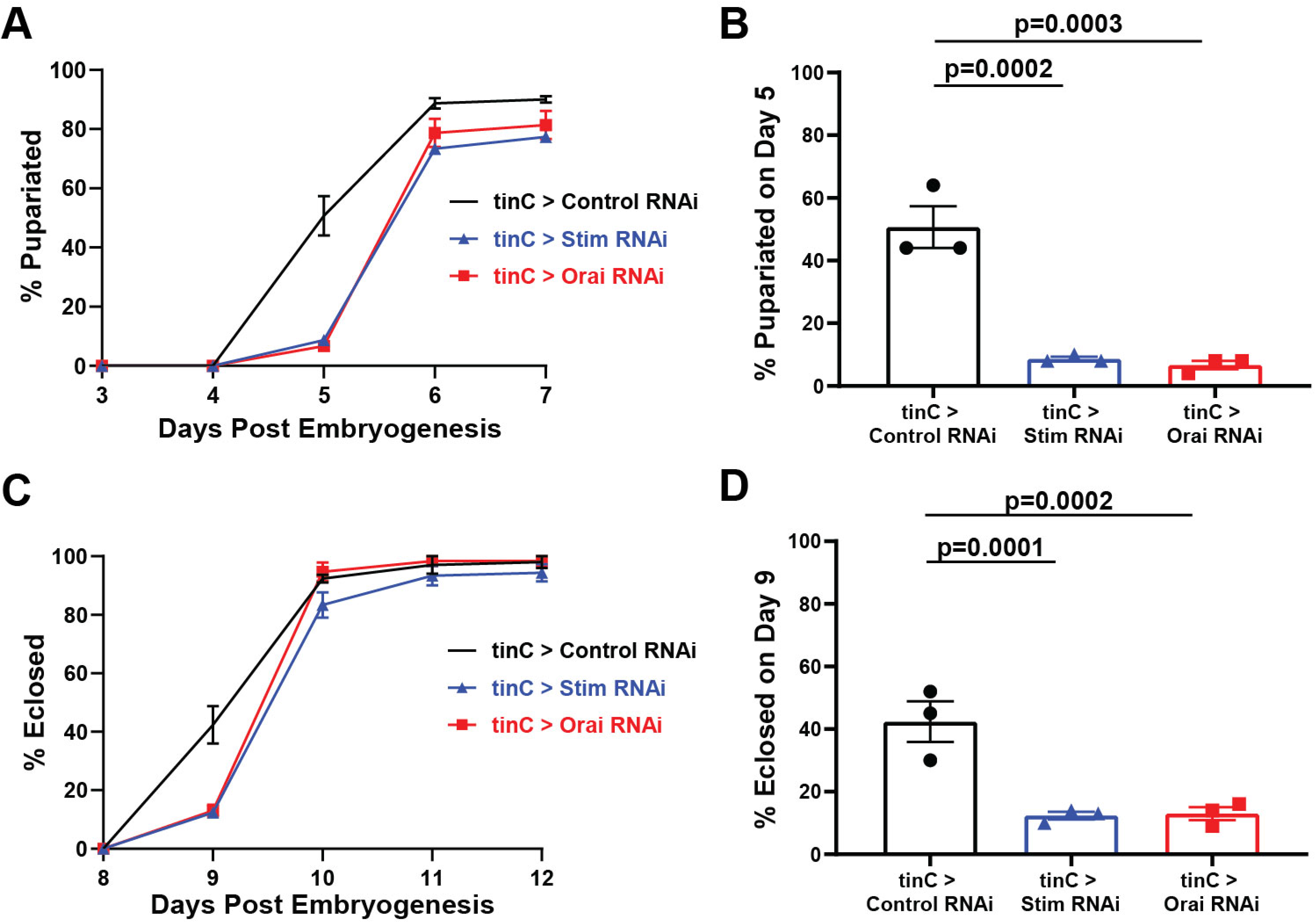
Heart specific *Stim* and *Orai* suppression delays post-embryonic animal development. **A.** Plot of the percent of larvae that pupariated on each of the indicated days post-embryogenesis for *tinC-GAL4* driven *Stim*, *Orai*, and non-targeting control RNAi. Data are mean ± SEM from three independent experiments, with 25-50 animals per experimental group. **B.** Comparison of percent pupariated on day 5 post-embryogenesis for *tinC-GAL4* driven *Stim*, *Orai,* and non-targeting control RNAi from three independent replicates (p-values calculated from One-way ANOVA with Turkey’s Multiple Comparisons Test). **C.** Plot of the percent of pupae that eclosed on each of the indicated days post-embryogenesis for *tinC-GAL4* driven *Stim*, *Orai,* and non-targeting control RNAi. Data are mean ± SEM from three independent experiments, with 25-50 animals per experimental group. **D.** Comparison of percent eclosed on day 9 post-embryogenesis for *tinC-GAL4* driven *Stim*, *Orai* and non-targeting control RNAi from three independent replicates (p-values calculated from One-way ANOVA with Turkey’s Multiple Comparisons Test).

We next analyzed adult lifespan to determine whether heart dysfunction resulting from *Stim* and *Orai* suppression affects adult viability. Males with *tinC-GAL4* driven *Orai* RNAi died significantly earlier than those with nontargeting control RNAi, as indicated by a statistically significant reduction of approximately 13 days in average median lifespan for *Orai* RNAi versus controls (Figure 6A and B). Animals with *tinC-GAL4* driven *Stim* RNAi also exhibited an approximately 11 day reduction in average median life span compared to controls, though this difference was not statistically significant (Figure 6A and B). More profound effects on adult lifespan were observed with *4xhand-GAL4* driven *Stim* and *Orai* RNAi, as the average median lifespan of adult males was 11 ± 1.73 and 13 ± 2.65 days for *Stim* and *Orai* RNAi, respectively, compared to 43.64 ± 3.38 days for controls (Figure 6C-D; mean ± SEM). Thus, whereas *Stim* and *Orai* suppression driven by *tinC-GAL4* resulted in greater deficiencies in heart contractility, suppression driven by *4xhand-GAL4* resulted in greater reductions in adult lifespan. This difference may result from broader tissue distribution of *4xhand-GAL4* expression compared to *tinC-GAL4*, and thus *4xhand-GAL4* driven RNAi may have had effects on animal survival independent of heart function. Consistent with this, many *4xhand-GAL4* driven *Stim* and *Orai* RNAi adults exhibited severe wing damage, including blistering and tearing of large sections of the wings (Figure 6E). This is likely due to *4xhand-GAL4* driven RNAi expression in wing hearts, which are muscles of cardiac origin that circulate hemolymph throughout the wings (Togel et al., 2013; Togel et al., 2008). Interestingly, this suggests that SOCE is also required for proper wing heart function, though we did not investigate this possibility further. However, we did consider the possibility that wing damage may have contributed to the early lethality of the *4xhand-GAL4* driven RNAi animals by causing them to become stuck in the food or on the vial walls. To test this and more clearly determine whether lethality is attributable to dysfunction of the primary heart, we repeated adult survival experiments with *4xhand-GAL4* driven RNAi animals whose wings were removed at the time of eclosion. As shown in Figure 6F and G, wingless *4xhand-GAL4* driven *Stim* and *Orai* RNAi animals indeed survived longer than those with wings (compare to Figure 6C and D), but these animals still died significantly earlier than corresponding wingless controls. Furthermore, the median lifespan of wingless *4xhand-GAL4* driven *Stim* and *Orai* RNAi animals was shorter than controls by approximately 12 days, similar to the result with *tinC-GAL4* driven RNAi, suggesting consistent early lethality due to abnormal heart function. Collectively, these results suggest that heart dysfunction caused by SOCE suppression pathologically impairs developmental physiology and longevity in *Drosophila*.

**Figure 6:**
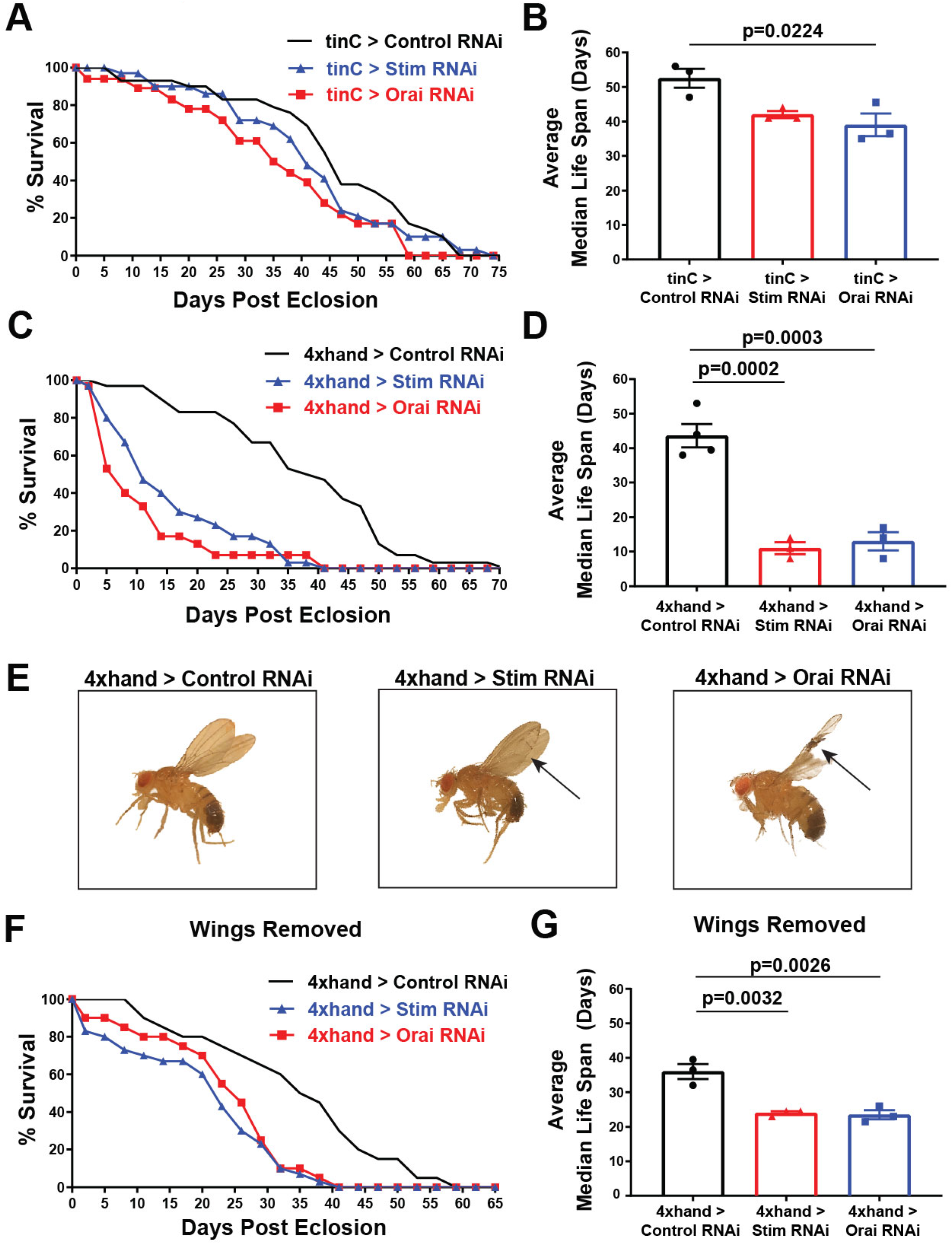
Heart specific *Stim* and *Orai* suppression results in early adult lethality. **A.** Representative survival curves in days post-eclosion for adult males with *tinC-GAL4* driven *Stim*, *Orai*, and non-targeting control RNAi (n = 30 animals per group). **B.** Plot of the average adult male median lifespan (MLS), calculated from three independent survival curves, for *tinC-GAL4* driven *Stim, Orai*, and non-targeting control RNAi. Error bars represent SEM, and p-values were calculated from One-way ANOVA with Turkey’s Multiple Comparisons Test **C.**Representative survival curves in days post-eclosion for adult males with *4xhand-GAL4* driven *Stim*, *Orai*, and non-targeting control RNAi (n = 30 animals per group). **D.** Plot of the average adult male median lifespan (MLS), calculated from three independent survival curves, for *4xhand-GAL4* driven *Stim, Orai,* and non-targeting control RNAi. Error bars represent SEM, and p-values were calculated from One-way ANOVA with Turkey’s Multiple Comparisons Test. **E.** Representative images of adult males with *4xhand-GAL4* driven *Stim*, *Orai*, and non-targeting control RNAi on the day of eclosion. Note the blistered wing on the *Stim* RNAi animal and the severely damaged and frayed wings on the *Orai* RNAi animal (indicated by arrows). **F.** Representative survival curves in days post-eclosion for adult males with *4xhand-GAL4* driven *Stim*, *Orai*, and non-targeting control RNAi, with their wings removed on day of eclosion (n = 30 animals per group). **G.** Plot of the average adult male median lifespan (MLS), calculated from three independent survival curves, for *4xhand-GAL4* driven *Stim*, *Orai*, and non-targeting control RNAi animals with their wings removed on the day of eclosion. Error bars represent SEM, and p-values were calculated from One-way ANOVA with Tukey’s Multiple Comparisons Test.

## DISCUSSION

The critical importance of proper Ca^2+^ homeostasis and transport in cardiomyocytes is accentuated by the fact that nearly all cardiomyopathies involve Ca^2+^ dysregulation. Clear understanding of the cellular and molecular mechanisms that regulate cardiomyocyte Ca^2+^ handling is therefore fundamental to unraveling the complexities of cardiac pathophysiology. Our results demonstrate that SOCE is an essential component of cardiomyocyte Ca^2+^ physiology in *Drosophila*, because suppression of the key molecular components of the SOCE pathway, *Stim* and *Orai,* resulted in contractile deficiencies that are highly consistent with dilated cardiomyopathy. In conjunction with contractility defects, SOCE suppressed hearts exhibited disrupted myofibril organization and density, a common finding in genetic cardiomyopathy models. Furthermore, our results demonstrate that reduced functional output of *Stim* and *Orai* suppressed hearts significantly impaired overall animal physiology and health, because these animals developed significantly more slowly and died earlier than controls. Importantly, our results are largely consistent with those from mice and zebrafish in which STIM1 or Orai1, respectively, were suppressed (Collins et al., 2014; Volkers et al., 2012), strongly suggesting that SOCE has essential, cardiomyocyte-specific functions that are highly conserved across animal species.

Our current study adds to previous findings showing cardiomyocyte-specific functions of STIM and Orai proteins in new and important ways. Most notably, our results are the first demonstration that suppression of both *Stim* and *Orai* results in nearly identical cardiac phenotypes in the same animal model. This suggests that these two proteins function together to regulate the same process, namely SOCE. This is significant, because previous studies in mammalian cardiomyocytes have implicated Orai1-independent targets of STIM1, such as canonical transient receptor potential (TrpC) channels (Ohba et al., 2009) and L-type Ca^2+^ channels (Correll et al., 2015), suggesting SOCE-independent functions of STIM1 in cardiomyocytes. While our results do not rule out additional functional partners of STIM and Orai, particularly in vertebrate species, they do strongly suggest that Ca^2+^ influx mediated specifically by SOCE is essential for proper cardiomyocyte physiology. This conclusion is further substantiated by the fact that *Drosophila* express single *Stim* and *Orai* isoforms. Thus, we can effectively target the SOCE pathway by suppressing only *Stim* or *Orai*, without possible compensation by other isoforms such as STIM2, Orai2, or Orai3. Moving forward, these advantages of the *Drosophila* model will allow us to investigate the functional roles of SOCE in cardiomyocytes using genetic tools that are largely unattainable with vertebrate models.

Despite several recent studies demonstrating that SOCE is required for normal, healthy heart function, the specific cellular processes that are regulated by SOCE in cardiomyocytes have not been clearly defined. Maintenance of SR Ca^2+^ stores is a seemingly likely function of SOCE that would broadly impact cardiomyocyte physiology. For example, reduced SR Ca^2+^ content in the absence of SOCE may reduce RyR-dependent Ca^2+^ release that drives contraction, and would be consistent with the severe systolic dysfunction that we and others have observed in STIM and Orai suppressed hearts. In this case, the myofibril disorganization observed in SOCE suppressed hearts may be the result of decompensation in response to reduced contractile force generation (Houser and Margulies, 2003). However, direct analyses of SR Ca^2+^ store content in STIM or Orai depleted cardiomyocytes have yielded conflicting results. For example, siRNA knockdown of STIM1 in freshly isolated neonatal rat ventricular myocytes led to a significant reduction of the caffeine-releasable SR Ca^2+^ pools (Voelkers et al., 2010), whereas SR Ca^2+^ store content was unchanged in adult cardiomyocytes isolated from STIM1 deleted mouse hearts compared to controls (Parks et al., 2016). These STIM1 deleted hearts also exhibited dilated cardiomyopathy, suggesting that SR Ca^2+^ store depletion may not be a direct cause of heart failure when SOCE is suppressed. However, an independent analysis of STIM1 deficiency-dependent dilated cardiomyopathy in mice demonstrated significantly upregulated expression of ER stress response factors (Collins et al., 2014), possibly reflecting mild but chronic Ca^2+^ store depletion that may be difficult to measure in acutely isolated cells. An important possibility is that reorganization of cytoarchitecture, including myofibril disarray, may be an effect of ER stress that impairs contractility and leads to heart failure.

It is also possible that SOCE has specific signaling functions in cardiomyocytes that are required for heart development or tissue homeostasis. The best characterized signaling function of SOCE across cell and tissue types involves calcineurin-mediated dephosphorylation and activation of transcription factors such as NFAT. Overwhelming evidence has established that upregulated calcineurin-NFAT signaling is an essential mechanism of cardiac hypertrophy pathogenesis (Tham et al., 2015), and SOCE upregulation also results in cardiac hypertrophy (Benard et al., 2016; Hulot et al., 2011; Parks et al., 2016) and enhanced calcineurin activity (Luo et al., 2012). Collectively, these findings suggest that calcineurin is a key cardiomyocyte target of SOCE in the pathological, upregulated state. However, whether calcineurin or other signaling factors are essential targets of SOCE in normal heart physiology has not been determined. Arguing against a signaling role of SOCE during heart development, STIM1 knockout mouse hearts do not exhibit altered phenotypes at early postnatal stages (Collins et al., 2014), and we also did not observe effects of heart-specific *Stim* or *Orai* suppression on *Drosophila* embryogenesis (data not shown). In addition, cardiomyopathy in STIM1 knockout mouse hearts was progressive and age-dependent (Collins et al., 2014), further suggesting that any signaling functions of SOCE are likely to be homeostatic rather than developmental. It is important to note that we first observed heart defects in larvae, early in the *Drosophila* lifecycle. This may also reflect homeostatic functions of SOCE, and the effects of SOCE suppression may manifest earlier in these animals due to the rapid lifecycle of *Drosophila* compared to vertebrate models. Clearly, much work is still required to identify the specific functional roles of SOCE in the healthy heart, and genetic tools combined with *in vivo* analyses of heart physiology and Ca^2+^ dynamics in *Drosophila* will allow us to address this in novel and significant ways.

Our study is the first to demonstrate the use of intravital fluorescence imaging of heart contractility in *Drosophila* larvae, an important lifecycle stage that involves rapid animal growth and high nutritional demands. Importantly, larval heart contractility has been difficult to analyze using other methods often employed for adults, such as OCT and direct imaging of dissected preparations. For example, the high fat content of larvae introduces light scattering that can obscure OCT imaging. Furthermore, the larval heart is extremely delicate and poorly attached to the larval body wall, making partial dissections that faithfully preserve heart function extremely challenging. Thus, intravital fluorescence imaging of larval hearts is a versatile and accessible method that is also amenable to longitudinal analysis. We first validated our intravital imaging approach by confirming that we were able to measure significantly increased ESD and EDD and decreased FS values in adult *Stim* suppressed compared to control hearts, similar to our results from the more established OCT method. However, it was notable that even in controls, ESD and EDD values were substantially larger with intravital imaging compared to OCT measurements. This is likely due to better spatial resolution of the heart walls with fluorescence imaging compared to OCT. Importantly, our fluorescence imaging measurements of adult heart ESD, EDD, and FS were very similar to those from the first study that developed the CM-tdTom model for intravital fluorescence imaging (Klassen et al., 2017), further validating the accuracy of our measurements.

Another notable finding of our study was that there were several key differences in the results of *Stim* and *Orai* RNAi driven with *tinC-GAL4* compared to *4xhand-GAL4*, both of which are described interchangeably in the literature as heart specific *GAL4* drivers. Most strikingly, *4xhand-GAL4* but not *tinC-GAL4* driven RNAi resulted in significant effects on adult wing structure and integrity that were likely caused by defective wing heart function, given that *hand* but not *tinman* (the genes from which the respective *GAL4* drivers are derived) is expressed in wing heart myocytes (Togel et al., 2013; Togel et al., 2008). Importantly, we found that the wing defects caused by *4xhand-GAL4* driven *Stim* and *Orai* RNAi significantly shortened animal lifespan, independently of effects on contractility of the primary adult heart. This illustrates that caution is warranted when attributing phenotypes, such as premature lethality, to dysfunction of the primary *Drosophila* heart when using these *GAL4* drivers.

In conclusion, our results demonstrate that SOCE mediated by STIM and Orai is essential for proper function of the *Drosophila* heart, and add to a growing number of studies that collectively suggest that SOCE has highly conserved functional roles in cardiomyocytes across animal species. Paradoxically, it is still unknown whether loss of function mutations in *STIM1* or *Orai1* result in heart defects or cardiomyopathies in humans likely due to the fact that individuals homozygous for these mutations die from immunodeficiency in infancy or early childhood, before adverse heart phenotypes may fully manifest (Rosenberg et al., 2019). Therefore, animal model studies will continue to be vital to our understanding of how alterations to SOCE result in or contribute to devastating cardiomyopathies and heart failure.

## MATERIALS AND METHODS

### Fly Stocks

The following *Drosophila* stocks were obtained from the Bloomington *Drosophila* Stock Center: mCherry RNAi (35785; non-targeting control), GAL4 RNAi (35783; non-targeting control RNAi), *Stim* RNAi (27263), *Orai* RNAi (53333), and *act-GAL4* (3954). *tinC-GAL4* was obtained from Dr. Manfred Frausch (Friedrich Alexander University), and *4xhand-GAL4* was from Dr. Zhe Han (George Washington University School of Medicine). CM-Tdtom flies were obtained from Dr. Rolf Bodmer (Sanford Burnham Prebys Institute). Flies were maintained on standard cornmeal agar food, and all crosses were carried out at 25°C.

### Adult Survival

Virgin female *tinC-GAL4* or *4xhand-GAL4* flies were crossed with male RNAi animals, and progeny were raised to adulthood at 25°C to eclosion. On the day of eclosion, adult progeny were collected and separated based on sex into vials containing up to ten flies per vial, with a total of 20-30 flies per group. Flies were maintained at 25°C throughout the course of the experiments. Every three days, the vials were checked for dead animals and surviving flies were transferred into new vials with fresh food. For wingless experiments, wings were removed immediately following eclosion on the day of adult collection.

### Developmental Timing

Approximately 30-40 virgin female *tinC-GAL4* or *4xhand-GAL4* animals were mated with RNAi males for three days, at which time mated females were transferred into egg laying chambers that consisted of a 100ml plastic beaker with holes for air exchange affixed over a petri dish containing grape juice agar (Genesee Scientific). A dollop of yeast paste (active dry yeast mixed with water) was placed in the center of each grape juice agar plate to provide food. Animals were acclimated in the chambers for 24 hours, and then transferred to new plates for 4 hours at 25°C for timed egg laying. After removing the females, plates with eggs were incubated at 25°C for an additional 24 hours. Hatched larvae were then transferred to vials with standard fly food, with up to 30 larvae per vial, and maintained at 25°C through the course of the experiment. Vials were checked each day, and the numbers of newly formed pupae and eclosed adults were recorded.

### RNA Isolation and RT-qPCR

Twenty to 30 first instar larvae were collected in 20μl cold Trizol and manually crushed with a pestle. Additional Trizol was then added to a total volume of 500μl, 100μl chloroform was added, and the aqueous layer containing extracted RNA was isolated. Extracted RNA was further purified with the RNeasy kit (Qiagen) and converted to cDNA using an S1000 Thermo Cycler (BioRad) with high capacity cDNA Reverse Transcription kit (ThermoFisher). Real-time quantitative polymerase chain reaction (RT-qPCR) was performed on a StepOnePlus RT-qPCR machine (Applied Biosystems) with each reaction consisting of triplicate samples containing iTaq Universal Probes Supermix (BioRad), pre-validated 6-carboxyfluorescein (FAM)-labeled Taqman probes (Applied Biosystems) against *Stim*, *Orai*, and *RPL32* (housekeeping gene), and template cDNA diluted per manufacturer’s instructions. For quantification, triplicate cycle threshold (Ct) values were averaged and normalized to the *RPL32* Ct value to calculate ΔCt. The Δ(ΔCt) was determined by subtracting the control RNAi ΔCt value from the experimental ΔCt value, and fold changes expressed as 2^−Δ(ΔCt)^. Fold changes are expressed as a percentage of expression compared to non-targeting RNAi control.

### Heart Dissection, Staining, and Confocal Imaging

Hearts from third instar larvae of five-day old adults were dissected and fixed as previously described (Alayari et al., 2009). In brief, for adults the ventral abdomens and underlying tissues were removed to expose the contracting heart while bathed in oxygenated artificial *Drosophila* hemolymph (ADH; 108 mM NaCl, 5 mM KCl, 2 mM CaCl_2_, 8 mM MgCl_2_, 1 mM NaH_2_PO_4_, 4 mM NaHCO_3_, 10 mM sucrose, 5 mM trehalose, and 5 mM HEPES (pH 7.1). Hearts were then fully relaxed by exchange with fresh ADH containing 10 mM EGTA. Larvae were pinned to Sylgard coated dishes at their anterior and posterior, and a slit was cut along the ventral midline in the presence of oxygenated ADH. Lateral cuts were then made along the sides of the animals, and the resulting cuticle flaps were pinned to expose the internal organs. The gut was removed to expose the beating hearts, which were then relaxed with ADH containing 10 mM EGTA. Dissected adults and larvae were both fixed for 20 min at room temperature in PBS containing 4% paraformaldehyde. Following three 10 min washes in PBS containing 0.1% Triton X-100 (PBSTx) with gentle rotation, the samples were incubated in PBSTx containing 1.0 μM Alexa Fluor 488 Phalloidin (ThermoFisher) for one hour at room temperature with gentle shaking. Samples were again washed three times in PBSTx at room temperature, and mounted on glass slides and coverslips with Vectashield (Vectashield Laboratories) as described (Alayari et al., 2009). Samples were imaged with a Nikon A1R confocal microscope using 10X, 0.45 N.A. and 40X, 1.3 NA objectives. Phalloidin was excited with a 488 nm laser. Z-stacks at 1 μm intervals were collected and images are presented as maximum intensity projections encompassing the whole heart for larvae or the dorsal half of the heart for adults to avoid the ventral layer of skeletal muscle. Myofibril density measurements on adults and third instar larval hearts were taken from the A2 segment of the heart. The number of myofibrils per 75μm in larvae and 50μm in adults was calculated manually.

### Optical Coherence Tomography (OCT)

Adult heart contractility was analyzed using a custom-built OCT apparatus as previously described (Wolf et al., 2006). In brief, five-day old males were briefly anesthetized with CO_2_, embedded in a soft gel support, and allowed to fully awaken based on body movement. Animals were first imaged in B-mode in the longitudinal orientation to identify the A1 segment of the heart chamber. They were then imaged transversely in M-mode for 3 sec, and multiple M-modes were recorded for each fly. Animals were then re-imaged in B-mode to ensure proper orientation of the heart chamber. M-modes were processed in ImageJ by referencing to a 150 μm standard. EDD, ESD, and heart rate were calculated directly from the processed M-mode traces. FS was calculated as (EDD-ESD) / EDD × 100.

### Intravital Fluorescence Microscopy

Intravital fluorescence imaging of adult and third instar larval hearts was carried out using animals that express tdTomato under control of the cardiomyocyte-specific R94C02 enhancer element (Klassen et al., 2017) Five-day old adult males and third instar larvae were briefly anesthetized with CO_2_ for adults or 1 minute cold exposure at 4°C for larva. Animals were then adhered dorsal side down to a glass coverslip with Norland Optical Adhesive and cured with a 365nm 3-watt UV LED light source (brand) for 30 seconds. Animals were allowed to recover for 10 mins prior to imaging. The heart was imaged through the dorsal cuticle at a rate of 200 frames per second (fps) for ten seconds using a sCMOS camera (Hamamatsu) on a Nikon Ti2 inverted microscope controlled with Nikon Elements software. Excitation light at 550 nm was provided by a Spectra-X illuminator (Lumencor) and emission was collected through a 555-635 band-pass filter. The full-time lapse of images acquired at 200 fps were used to generate M-modes from the A1 segment of the heart chamber in adults and the A2 segment in third instar larva using ImageJ (NIH). EDD, ESD, and heart rate were calculated directly from the processed M-mode traces. FS was calculated as (EDD-ESD) / EDD × 100.

## ACKNOWLEDGEMENTS

This work was supported by funding from the Collaborative Health Initiative Research Program to C.E.P. and J.T.S. and from the National Institutes of Health to M.J.W.

## Notes

#### Summary of Updates

Additional figures and data have been added, including intravital imaging of Drosophila heart contractility.

## REFERENCES

Abraham, D.M., and Wolf, M.J. (2013). Disruption of sarcoendoplasmic reticulum calcium ATPase function in Drosophila leads to cardiac dysfunction. PloS one 8, e77785.

Alayari, N.N., Vogler, G., Taghli-Lamallem, O., Ocorr, K., Bodmer, R., and Cammarato, A. (2009). Fluorescent labeling of Drosophila heart structures. Journal of visualized experiments : JoVE.

Benard, L., Oh, J.G., Cacheux, M., Lee, A., Nonnenmacher, M., Matasic, D.S., Kohlbrenner, E., Kho, C., Pavoine, C., Hajjar, R.J., et al. (2016). Cardiac Stim1 Silencing Impairs Adaptive Hypertrophy and Promotes Heart Failure Through Inactivation of mTORC2/Akt Signaling. Circulation 133, 1458–1471; discussion 1471.

Bers, D.M. (2002). Cardiac excitation-contraction coupling. Nature 415, 198–205.

Collins, H.E., He, L., Zou, L., Qu, J., Zhou, L., Litovsky, S.H., Yang, Q., Young, M.E., Marchase, R.B., and Chatham, J.C. (2014). Stromal interaction molecule 1 is essential for normal cardiac homeostasis through modulation of ER and mitochondrial function. American journal of physiology Heart and circulatory physiology 306, H1231–1239.

Correll, R.N., Goonasekera, S.A., van Berlo, J.H., Burr, A.R., Accornero, F., Zhang, H., Makarewich, C.A., York, A.J., Sargent, M.A., Chen, X., et al. (2015). STIM1 elevation in the heart results in aberrant Ca(2)(+) handling and cardiomyopathy. Journal of molecular and cellular cardiology 87, 38–47.

Garfinkel, A.C., Seidman, J.G., and Seidman, C.E. (2018). Genetic Pathogenesis of Hypertrophic and Dilated Cardiomyopathy. Heart failure clinics 14, 139–146.

Horton, J.S., Buckley, C.L., Alvarez, E.M., Schorlemmer, A., and Stokes, A.J. (2014). The calcium release-activated calcium channel Orai1 represents a crucial component in hypertrophic compensation and the development of dilated cardiomyopathy. Channels (Austin, Tex) 8, 35–48.

Houser, S.R., and Margulies, K.B. (2003). Is depressed myocyte contractility centrally involved in heart failure? Circulation research 92, 350–358.

Hulot, J.S., Fauconnier, J., Ramanujam, D., Chaanine, A., Aubart, F., Sassi, Y., Merkle, S., Cazorla, O., Ouille, A., Dupuis, M., et al. (2011). Critical role for stromal interaction molecule 1 in cardiac hypertrophy. Circulation 124, 796–805.

Klassen, M.P., Peters, C.J., Zhou, S., Williams, H.H., Jan, L.Y., and Jan, Y.N. (2017). Age-dependent diastolic heart failure in an in vivo Drosophila model. eLife 6.

Kranias, E., and Bers, D. (2007). Calcium and Cardiomyopathies, Vol 45.

Limas, C.J., Olivari, M.T., Goldenberg, I.F., Levine, T.B., Benditt, D.G., and Simon, A. (1987). Calcium uptake by cardiac sarcoplasmic reticulum in human dilated cardiomyopathy. Cardiovascular research 21, 601–605.

Lin, N., Badie, N., Yu, L., Abraham, D., Cheng, H., Bursac, N., Rockman, H.A., and Wolf, M.J. (2011). A method to measure myocardial calcium handling in adult Drosophila. Circulation research 108, 1306–1315.

Luo, X., Hojayev, B., Jiang, N., Wang, Z.V., Tandan, S., Rakalin, A., Rothermel, B.A., Gillette, T.G., and Hill, J.A. (2012). STIM1-dependent store-operated Ca(2)(+) entry is required for pathological cardiac hypertrophy. Journal of molecular and cellular cardiology 52, 136–147.

MacLeod, K.T. (2016). Recent advances in understanding cardiac contractility in health and disease. F1000Research 5.

McKenna, W.J., Maron, B.J., and Thiene, G. (2017). Classification, Epidemiology, and Global Burden of Cardiomyopathies. Circulation research 121, 722–730.

Molkentin, J.D., Lu, J.R., Antos, C.L., Markham, B., Richardson, J., Robbins, J., Grant, S.R., and Olson, E.N. (1998). A calcineurin-dependent transcriptional pathway for cardiac hypertrophy. Cell 93, 215–228.

Ocorr, K., Perrin, L., Lim, H.Y., Qian, L., Wu, X., and Bodmer, R. (2007). Genetic control of heart function and aging in Drosophila. Trends in cardiovascular medicine 17, 177–182.

Ohba, T., Watanabe, H., Murakami, M., Sato, T., Ono, K., and Ito, H. (2009). Essential role of STIM1 in the development of cardiomyocyte hypertrophy. Biochemical and biophysical research communications 389, 172–176.

Parks, C., Alam, M.A., Sullivan, R., and Mancarella, S. (2016). STIM1-dependent Ca(2+) microdomains are required for myofilament remodeling and signaling in the heart. Scientific reports 6, 25372.

Pathak, T., Trivedi, D., and Hasan, G. (2017). CRISPR-Cas-Induced Mutants Identify a Requirement for dSTIM in Larval Dopaminergic Cells of Drosophila melanogaster. G3 (Bethesda, Md) 7, 923–933.

Piazza, N., and Wessells, R.J. (2011). Drosophila models of cardiac disease. Progress in molecular biology and translational science 100, 155–210.

Putney, J.W. (2011). The physiological function of store-operated calcium entry. Neurochemical research 36, 1157–1165.

Putney, J.W. (2018). Forms and functions of store-operated calcium entry mediators, STIM and Orai. Adv Biol Regul 68, 88–96.

Rosenberg, P., Katz, D., and Bryson, V. (2019). SOCE and STIM1 signaling in the heart: Timing and location matter. Cell calcium 77, 20–28.

Rotstein, B., and Paululat, A. (2016). On the Morphology of the Drosophila Heart. Journal of Cardiovascular Development and Disease 3, 15.

Schulz, R.A., and Yutzey, K.E. (2004). Calcineurin signaling and NFAT activation in cardiovascular and skeletal muscle development. Developmental biology 266, 1–16.

Smyth, J.T., Hwang, S.Y., Tomita, T., DeHaven, W.I., Mercer, J.C., and Putney, J.W. (2010). Activation and regulation of store-operated calcium entry. Journal of cellular and molecular medicine 14, 2337–2349.

Sussman, M.A., Welch, S., Cambon, N., Klevitsky, R., Hewett, T.E., Price, R., Witt, S.A., and Kimball, T.R. (1998). Myofibril degeneration caused by tropomodulin overexpression leads to dilated cardiomyopathy in juvenile mice. The Journal of clinical investigation 101, 51–61.

Tham, Y.K., Bernardo, B.C., Ooi, J.Y., Weeks, K.L., and McMullen, J.R. (2015). Pathophysiology of cardiac hypertrophy and heart failure: signaling pathways and novel therapeutic targets. Archives of toxicology 89, 1401–1438.

Togel, M., Meyer, H., Lehmacher, C., Heinisch, J.J., Pass, G., and Paululat, A. (2013). The bHLH transcription factor hand is required for proper wing heart formation in Drosophila. Developmental biology 381, 446–459.

Togel, M., Pass, G., and Paululat, A. (2008). The Drosophila wing hearts originate from pericardial cells and are essential for wing maturation. Developmental biology 318, 29–37.

Vega, R.B., Bassel-Duby, R., and Olson, E.N. (2003). Control of cardiac growth and function by calcineurin signaling. The Journal of biological chemistry 278, 36981–36984.

Voelkers, M., Salz, M., Herzog, N., Frank, D., Dolatabadi, N., Frey, N., Gude, N., Friedrich, O., Koch, W.J., Katus, H.A., et al. (2010). Orai1 and Stim1 regulate normal and hypertrophic growth in cardiomyocytes. Journal of molecular and cellular cardiology 48, 1329–1334.

Volkers, M., Dolatabadi, N., Gude, N., Most, P., Sussman, M.A., and Hassel, D. (2012). Orai1 deficiency leads to heart failure and skeletal myopathy in zebrafish. Journal of cell science 125, 287–294.

Wilkins, B.J., Dai, Y.S., Bueno, O.F., Parsons, S.A., Xu, J., Plank, D.M., Jones, F., Kimball, T.R., and Molkentin, J.D. (2004). Calcineurin/NFAT coupling participates in pathological, but not physiological, cardiac hypertrophy. Circulation research 94, 110–118.

Wolf, M.J., Amrein, H., Izatt, J.A., Choma, M.A., Reedy, M.C., and Rockman, H.A. (2006). Drosophila as a model for the identification of genes causing adult human heart disease. Proceedings of the National Academy of Sciences of the United States of America 103, 1394–1399.

