## Supplemental Figures and Legends for "Suppression of store-operated calcium entry causes dilated cardiomyopathy of the *Drosophila* heart"

#### Supplementary Materials

### Suplemental Figure 1

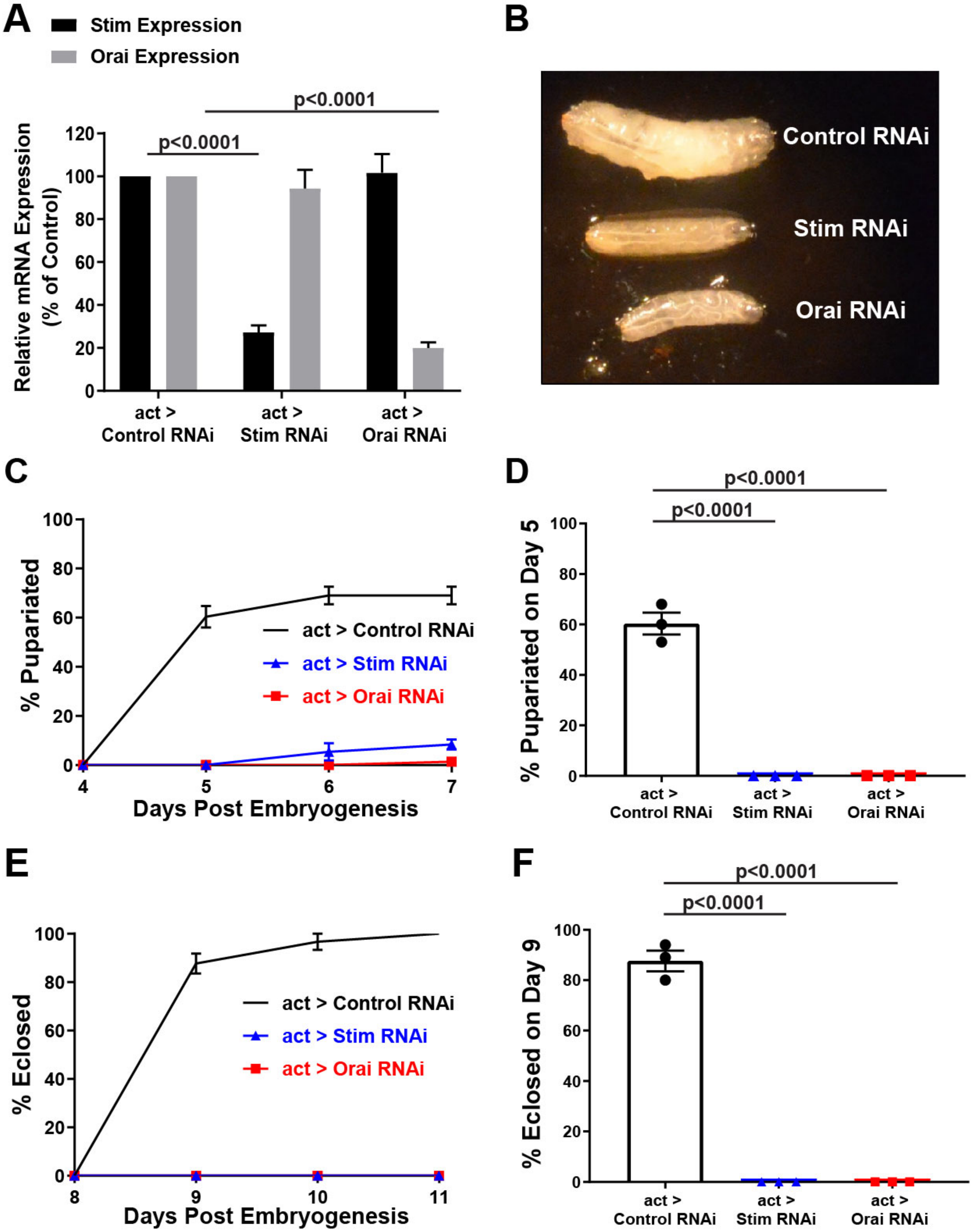

**Supplementary Figure S1.** ***Stim*** **and** ***Orai*** **RNAi are efficient and specific**

**A.** RT-qPCR analysis of relative *Stim* and *Orai* mRNA expression levels from first instar larvae with *act-GAL4* driven non-targeting control, *Stim*, and *Orai* RNAi. Data are represented as percent of non-targeting control RNAi (mean ± SEM from three independent replicates and p-values were calculated from Two-way ANOVA with Tukey's Multiple Comparison). **B.** Representative images of third instar larvae with *act-GAL4* driven non-targeting control, *Stim*, and *Orai* RNAi. Note the significantly reduced size of *Stim* and *Orai* RNAi animals compared to controls. **C.** Plot of the percent of larvae that pupariated on each of the indicated days post-embryogenesis for *act-GAL4* driven *Stim*, *Orai*, and non-targeting control RNAi. Data are mean ± SEM from three independent experiments, with 25-50 animals per experimental group. **D.** Comparison of percent pupariated on day 5 post-embryogenesis for *act-GAL4* driven *Stim*, *Orai,* and non-targeting control RNAi from three independent replicates (p-values calculated from One-way ANOVA with Turkey's Multiple Comparisons Test). **E.** Plot of the percent of pupae that eclosed on each of the indicated days post-embryogenesis for *act-GAL4* driven *Stim*, *Orai,* and non-targeting control RNAi. Data are mean ± SEM from three independent experiments, with 25-50 animals per experimental group. **F.** Comparison of percent eclosed on day 9 post-embryogenesis for *act-GAL4* driven *Stim*, *Orai* and non-targeting control RNAi from three independent replicates (p-values calculated from One-way ANOVA with Turkey's Multiple Comparisons Test).

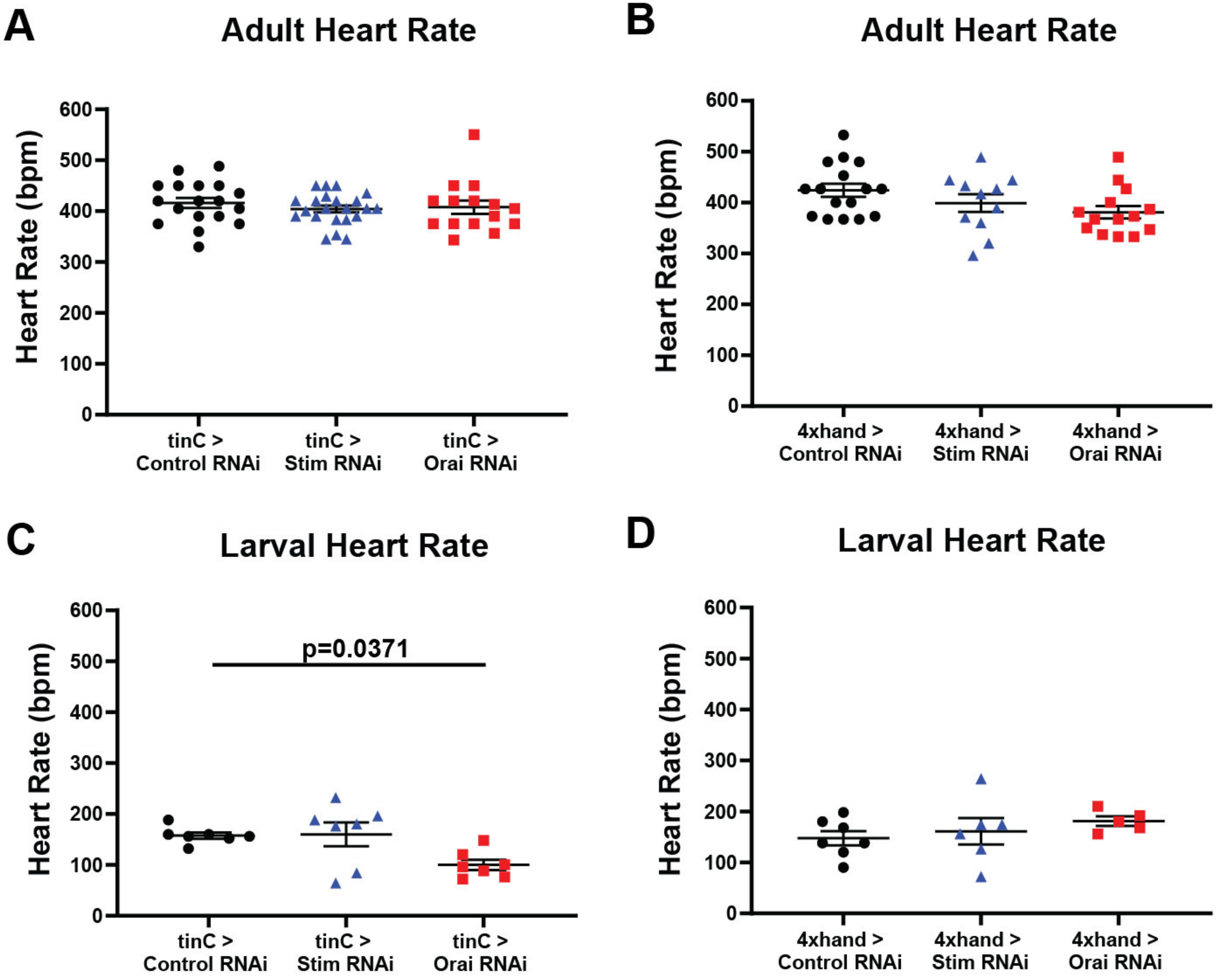


Supplemental Figure 2

__Supplementary Figure S2.__ **Heart specific suppression of *Stim* and *Orai* does not affect adult or larval heart rate**

Adult heart rate calculated as beats per min (bpm) from OCT imaging for *tinC-GAL4* (**A**) and *4xhand-GAL4* (**B**) driven non-targeting control, *Stim*, and *Orai* RNAi animals. Each symbol represents a single animal measurement; results were not significantly different (One-way ANOVA with Tukey's Multiple Comparisons Test). Third instar larval heart rate calculated as beats per min (bpm) from intravital fluorescence imaging for *tinC-GAL4* (**C**) and *4xhand-GAL4* (**D**) driven non-targeting control, *Stim* and *Orai* RNAi. Each symbol represents a single animal measurement; results were not significantly different except where indicated (p-value calculated from One-way ANOVA with Tukey's Multiple Comparison).

### Supplemental Figure 3

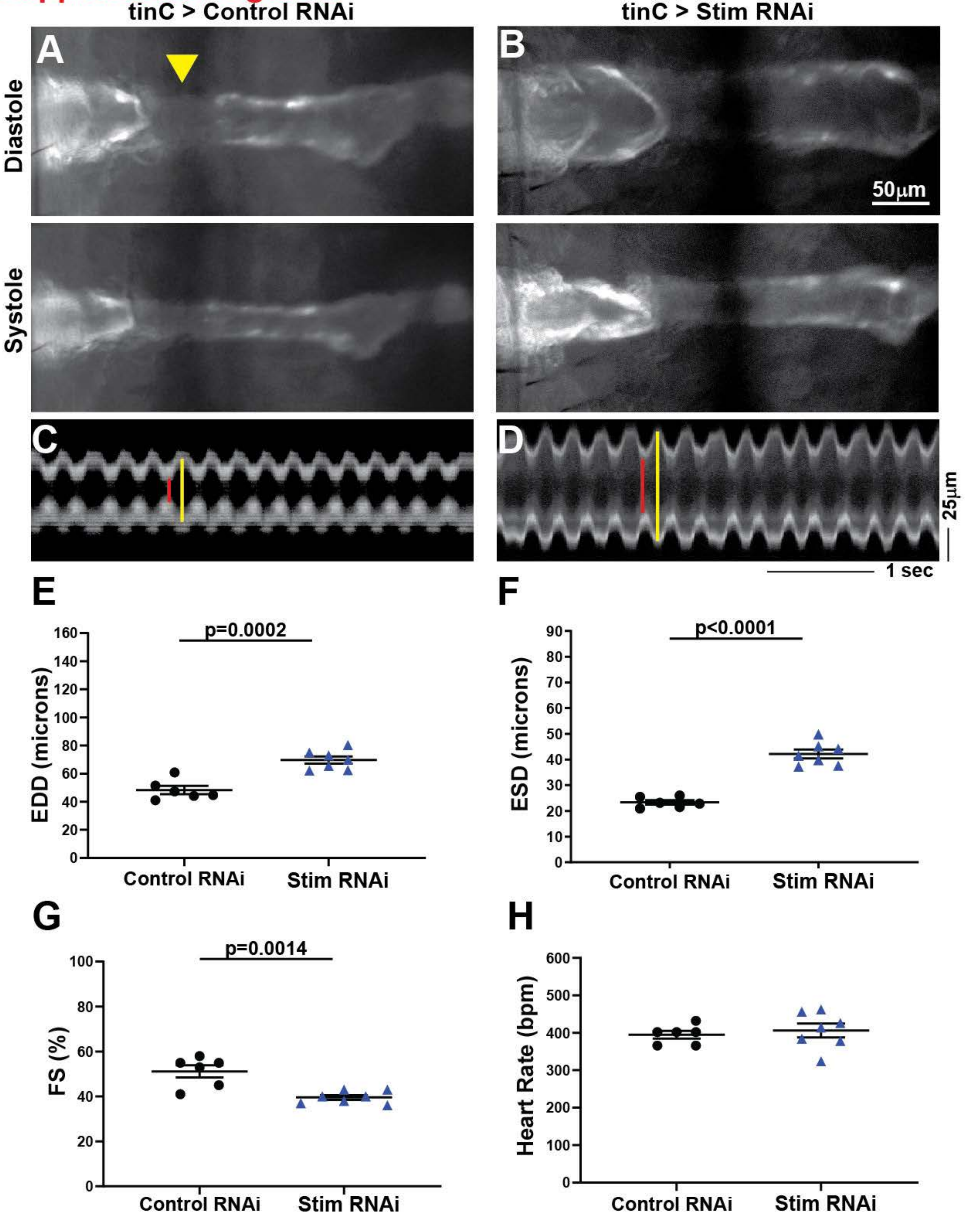

**Supplementary Figure S3: Intravital fluorescence imaging confirms dilated cardiomyopathy in adult *Stim* suppressed hearts**

**A-B.** Representative longitudinal B-mode images of intravital fluorescence imaging of R94C02-tdTom expressing five-day old adult male hearts with *tinC-GAL4* driven non-targeting control and *Stim* RNAi during diastole (upper panels) and systole (lower panels). Arrowhead points to region that is obscured by a dark abdominal stripe on the cuticle of the animal. **C-D.** Representative M-mode images from *tinC-GAL4* driven non-targeting control and *Stim* RNAi hearts, with red lines depicting systole and yellow lines diastole. EDD (**E**), ESD (**F**), FS (**G**), and heart rate (**H**) were calculated from M-mode recordings of five-day old males, and each symbol represents a measurement from a single animal. Bars indicate mean ± SEM, and p-values were calculated from unpaired t-tests.

#### SUPPLEMENTAL VIDEOS

**Supplemental Video S1. Full intravital imaging timelapse of the adult control heart shown in Figure S3.**

Images were acquired at a rate of 200 frames per second, and video playback is slowed to 100 frames per second to allow better visualization of contractions.

**Supplemental Video S2. Full intravital imaging timelapse of the adult *Stim* RNAi heart shown in Figure S3.**

Images were acquired at a rate of 200 frames per second, and video playback is slowed to 100 frames per second to allow better visualization of contractions.

**Supplemental Video S3. Full intravital imaging timelapse of the larval control heart shown in Figure 2A.**

Images were acquired at a rate of 200 frames per second, and video playback is at full frame-rate.

**Supplemental Video S4. Full intravital imaging timelapse of the larval *Stim* RNAi heart shown in Figure 2B.**

Images were acquired at a rate of 200 frames per second, and video playback is at full frame-rate.
